# Blood pressure variability from intra-arterial recordings in humans

**DOI:** 10.1101/775866

**Authors:** Farhan Adam Mukadam, Naveen Gangadharan, Bowya Baskaran, S Baskaran, Kandasamy Subramani, Syrpailyne Wankhar, Suresh Devasahayam, Sathya Subramani

## Abstract

Systolic and diastolic blood pressures are reported as single point values by the non-invasive techniques used in clinical practice, while, in fact, they are highly varying signals. The objective of this study was to document the magnitude of variation of systolic and diastolic pressures over a few minutes by analysing intra-arterial pressure recordings made in 51 haemodynamically stable patients in an intensive care unit. Intra-arterial pressure data were acquired by a validated data acquisition system. Fast-Flush test was performed and the dynamic characteristics of the catheter transducer system namely natural frequency and damping co-efficient were calculated. Only those recordings with acceptable dynamic characteristics were included in the analysis. Power spectral calculation using the Discrete Fourier transform (DFT) of the pressure recording revealed two frequency peaks below the peak at heart rate. The lower and higher frequency peaks below the heart rate peak are referred to as Mayer and Traube waves in this study. Mayer wave peaks were observed in DFT spectra of 49 out of 51 patients. The Mayer wave frequency peaks ranged between 0.045 Hz to 0.065 Hz in 41 out of 51 patients. The frequency of Traube waves or the respiratory variations was more than 0.14 Hz. Three categories of systolic and diastolic pressure variabilities namely beat-to-beat variability, Respiratory variability (Traube wave amplitude) and Total magnitude of variation are reported for all 51 patients. The mean systolic and diastolic pressure variations (in a period of about 10 minutes) in the study sample were 21 ± 9 mm Hg and 14 ± 5 mm Hg respectively. Given the magnitude of systolic and diastolic pressure variations over a few minutes, the validity of reporting single point values for these pressures and using single point cut-offs for diagnosis and treatment of hypertension must be re-evaluated.

## Introduction

Blood pressure and its measurement are of vital importance in clinical evaluation of cardiovascular status. The two most popular non-invasive methods of estimation of blood pressure are the manual method, based on Korotkoff sounds and the automated method based on cuff pressure oscillations. Both these methods report single point values for systolic and diastolic pressures. However, it is common knowledge that systolic and diastolic blood pressures vary at multiple frequencies, with wavelengths of the order of seconds to minutes (very short term Blood Pressure Variability, (BPV)), hours (24-hour BPV, short-term BPV), weeks (mid-term BPV) or even seasons (long-term BPV) [1]. In fact, no two consecutive pressure cycles have the same systolic and diastolic blood pressures and the concept of beat to beat variability is well-established. Such variations in systolic and diastolic pressures can be appreciated best in an intra-arterial recording using a fluid-filled catheter connected to a pressure transducer, which being a direct measurement of the pressure in the artery may be considered the gold standard for BP measurements [2], provided care is taken to avoid measurement errors due to air-bubbles, blood clots and compliance of the tube in the catheter system.

Intra-arterial BP is an invasive method and it is not possible to perform measurements with invasive techniques in routine clinical practice. There are non-invasive methods which claim to provide real time arterial pressure measurements. These methods are the volume-clamp method using a finger cuff and radial artery applanation tonometry. Some of these techniques and their decidedly limited usefulness are reported by [3-5].

Given the variability of systolic (SBP) and diastolic (DBP) blood pressures, the usefulness and credibility of making single point estimates of the two pressures with the non-invasive methods used in routine clinical practice needs re-evaluation.

Determining the variability of SBP and DBP is of crucial importance in understanding cardiovascular health. The role of blood pressure variations in target organ damage has been documented extensively [6-8]. Short term variability in both systolic and diastolic pressures is purported to be an independent risk factor for cardiovascular events or stroke [9]. It is also reported that antihypertensive drugs may either augment or reduce the variations in pressure. Atenelol is associated with a higher BPV, and does not reduce the incidence of stroke, even though it reduces mean systolic blood pressure. Addition of diuretics and calcium channel blockers to atenelol help in reducing risk of stroke and such reduction in stroke incidence is associated with a reduction in blood pressure variability [9].

This study reports the amplitude and nature of variations in SBP and DBP over a period of few minutes from intra-arterial recordings made in humans. Currently published information about blood pressure variation is limited and several published reports show poorly controlled dynamic characteristics of the fluid-filled catheter pressure measurement system making their data of limited value. Unless the pressure measurement system has a sufficient frequency bandwidth and appropriate damping, the measured waveform can be significantly erroneous.

It is desirable to have normative data for blood pressure variability for various categories of the population. As a first step, we have recorded intra-arterial blood pressure data from 51 patients in a surgical intensive care unit. The objective was to document the degree of variability seen in individuals, even though it is not a normal/control population, to illuminate the approximation in current non-invasive methods in routine clinical practice that report a single point value for SBP and DBP.

## Methods

Ethical and research aspects of this study were approved by the institutional review board (IRB). Conscious patients in the surgical ICU, with arterial pressure cannulae placed as standard of care were recruited in the study, after obtaining informed consent, when their blood pressure was stable, and they were not on vasoconstrictor or inotrope support. The transducer by design is positioned between the intra-arterial catheter and a pressurized fluid reservoir containing sterile perfusion fluid. The pressure at the transducer would be a value between the pressure in the artery and the pressure in the reservoir. The pressurized reservoir is held at a pressure of about 300 to 400 mm Hg. The resistance between the reservoir and the transducer is much higher than the resistance between the intra-arterial catheter and the transducer. Therefore, the pressure at the transducer is closer to the intra-arterial pressure and exceeds it by a negligible amount.

The higher pressure at the catheter ensures that blood does not enter the tubing connecting the catheter to the transducer. The pressure transducer output was connected to a data acquisition system (CMCdaq, a validated data recorder used in several labs in our institution) and data were acquired at a sampling frequency of 1KHz The pressure transducer was positioned at the level of intersection of the 4th intercostal space with the mid-axillary line (phlebostatic axis) and the three-way valve was opened to air for zeroing with respect to atmospheric pressure. The three-way valve was then switched to connect the transducer to the intra-arterial catheter and recording was begun.

Gardner [10] recommends that the natural frequency and damping coefficient of the pressure recording system be reported in all cases where accuracy of systolic pressure measurement is critical. These two parameters were assessed with a fast-flush test where a high-pressure pulse is applied to the transducer (with a pull of the fast-flush valve provided in the disposable pressure transducer, opening it to the high-pressure bag), followed by a quick release, allowing the catheter-transducer system to oscillate at its natural frequency on termination of the applied high-pressure plateau.

The natural frequency (ω_n_) and the damping coefficient (*ζ*) of the system were calculated from the oscillating waves at the end of the pressure pulse (Figure 1). Gardner [10] reports combinations of the two parameters (ω_n_ and *ζ*) which are adequate for accurate pressure recording in the form of a plot (see figure 2). The inner shaded area in figure 2, defines the boundaries for heart rates about 118 beats per minute (bpm) and the outer envelope defines the boundaries for a heart rate of 95 bpm.

**Figure 1:**
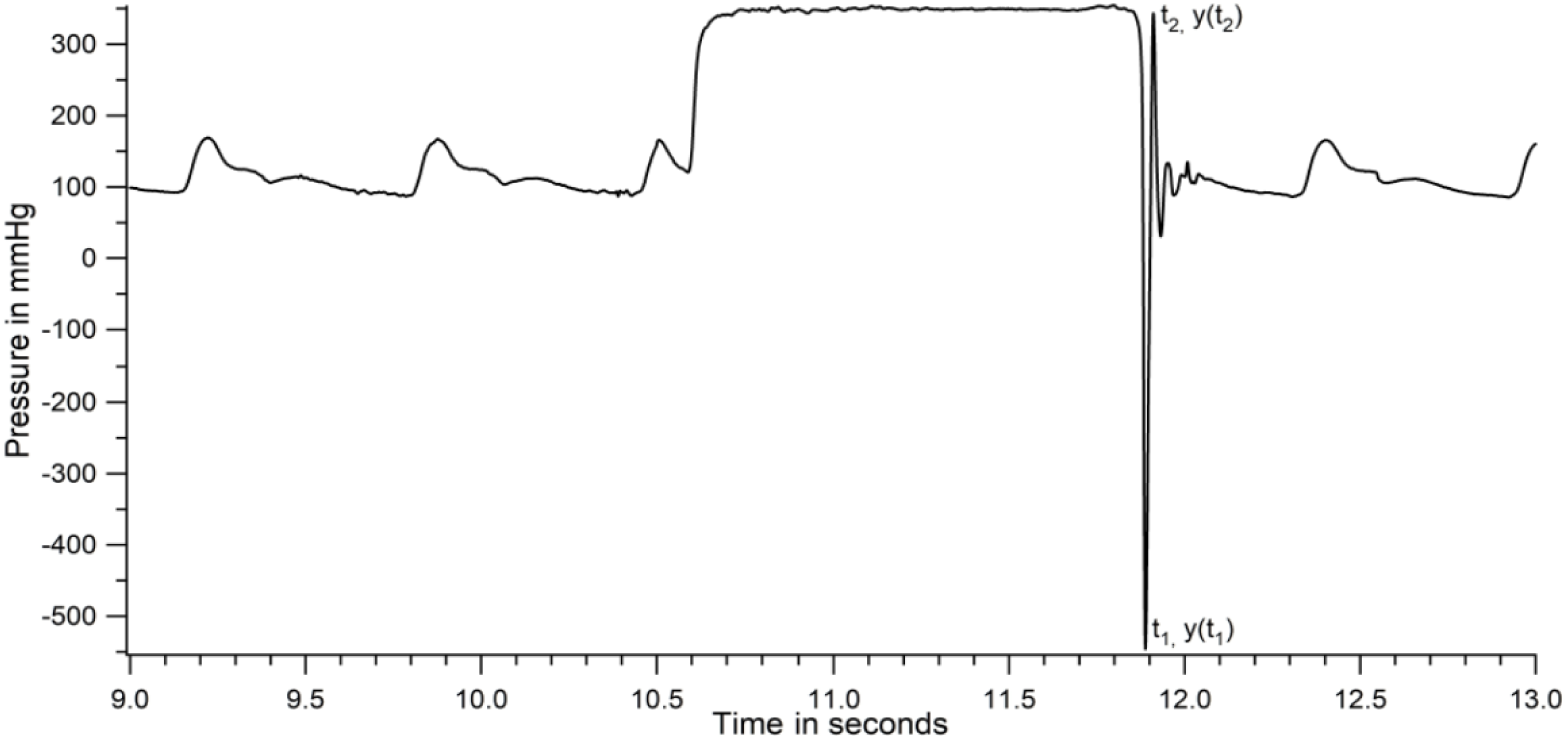
Dynamic response of catheter transducer system during a fast-flush test in a patient

**Figure-2:**
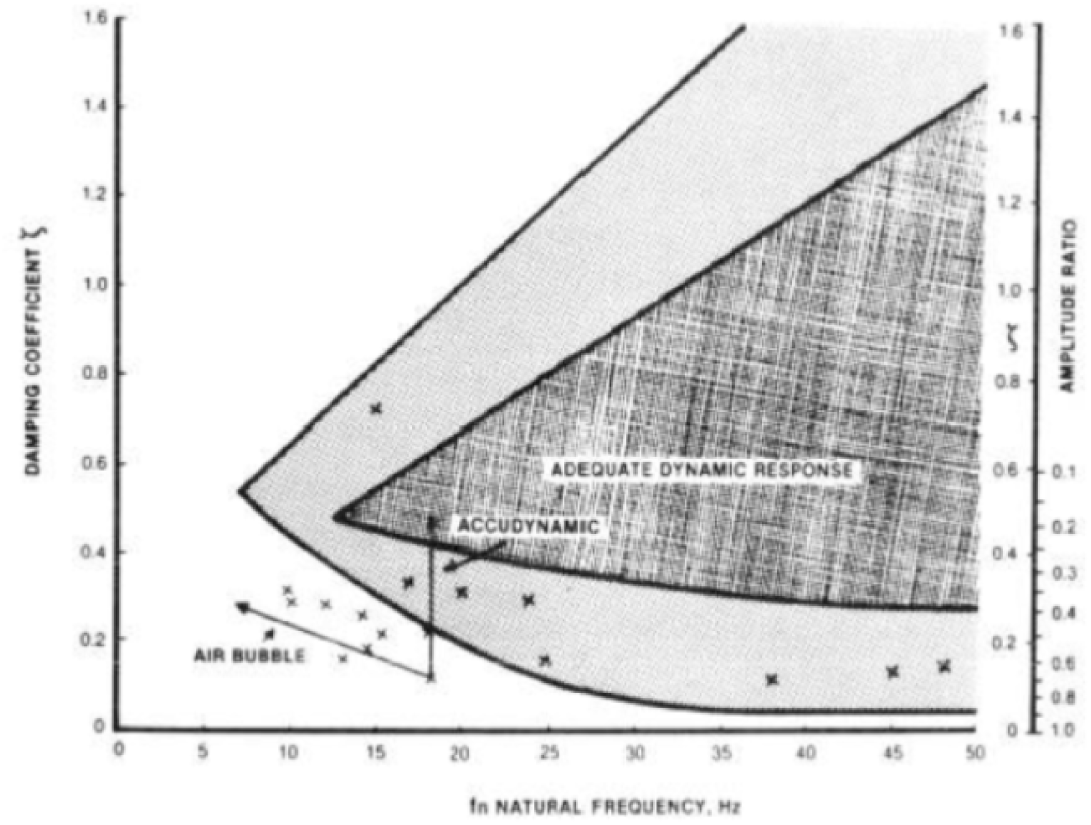
Boundary conditions for determining adequacy of the dynamic characteristics of the catheter-transducer system -Reproduced with permission from Gardner (1981)

From the above discussion, the higher the heart rate, the higher should be the natural frequency of the catheter-transducer system and narrower the range of acceptable damping coefficients. For want of better assessment of the adequacy of the dynamic response of the catheter system, we have obtained the two parameters for every patient recording, plotted them against each other and overlaid them on the Gardner plot (Figure 3).

**Figure-3.**
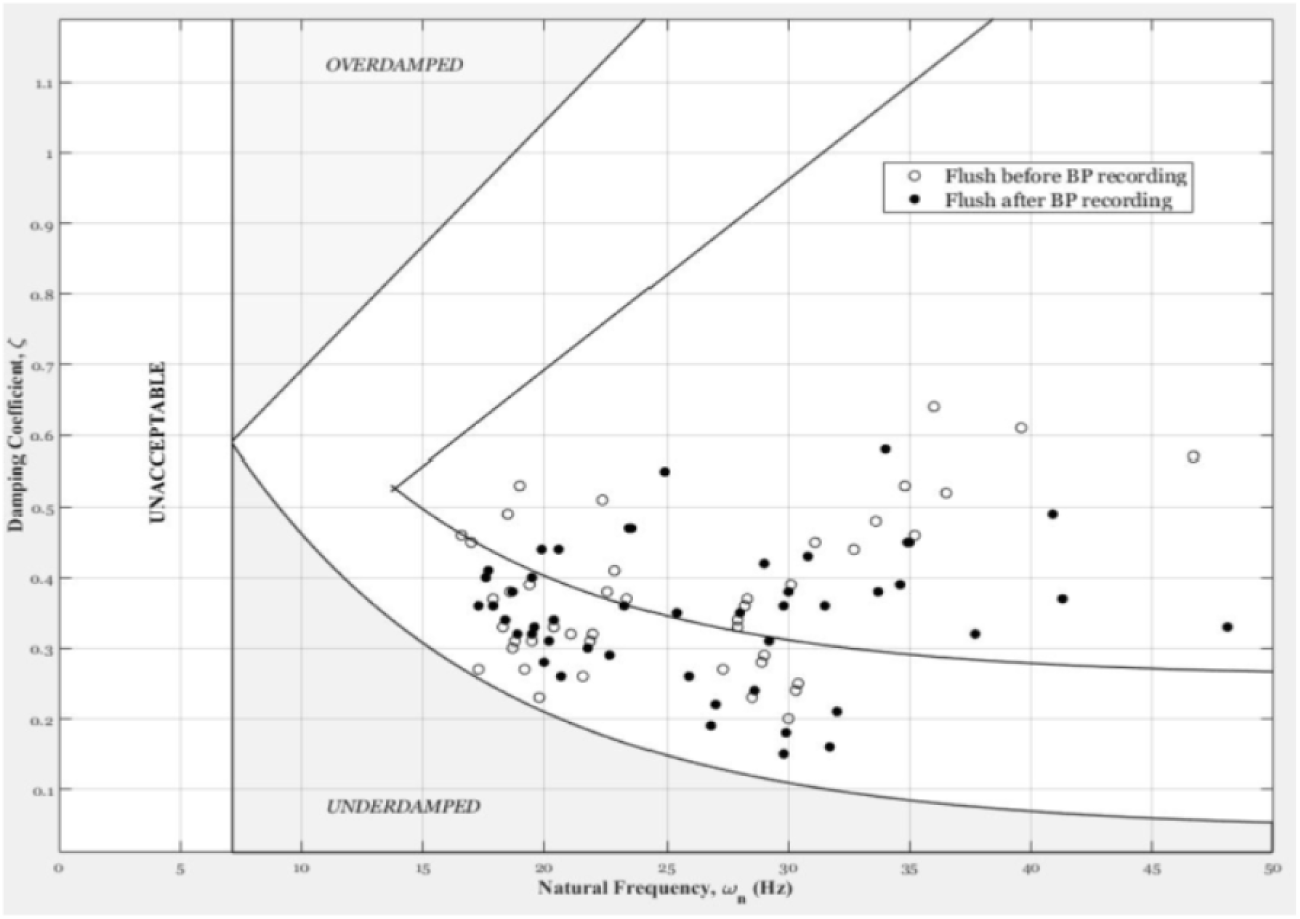
Plot of natural frequency versus damping coefficient for flush test data from 51 patients superimposed over the reference Gardner plot

The natural frequency of the recording system and the damping coefficient were calculated as described below. The time and amplitude of at least two extrema (minimum and/or maximum) are measured. Assuming that the flush test, or step response, can be approximated by a simple second order system response, the natural frequency and the damping coefficient were calculated using standard procedures. Briefly stated, the period of oscillation yields the damped frequency of the system, and the rate of decline of the amplitude swing yields the damping coefficient; the damped frequency corrected with the damping coefficient gives the natural frequency of the system. It is assumed in our calculations that the system oscillates around the mean arterial pressure of the pulses around the flush test.

The damping coefficient for flush test from each patient was plotted against the natural frequency (Figure 3). If the point fell within the acceptable boundaries shown in Figure 3, the patient data was included in the analysis. (The boundary lines of the Gardner plot were reproduced for Figure 3 by superimposition on the original plot).

## Results

Intra-arterial pressure data from 51 recordings with acceptable flush test criteria were analysed. Systolic and diastolic pressures in all individuals varied considerably during the duration of recording. Comprehensive analysis of one dataset is first presented to demonstrate the methodology of analysis. Figure 4 is the raw data obtained from a patient. Figure 5 shows a section of figure 4, expanded for clarity.

**Figure-4:**
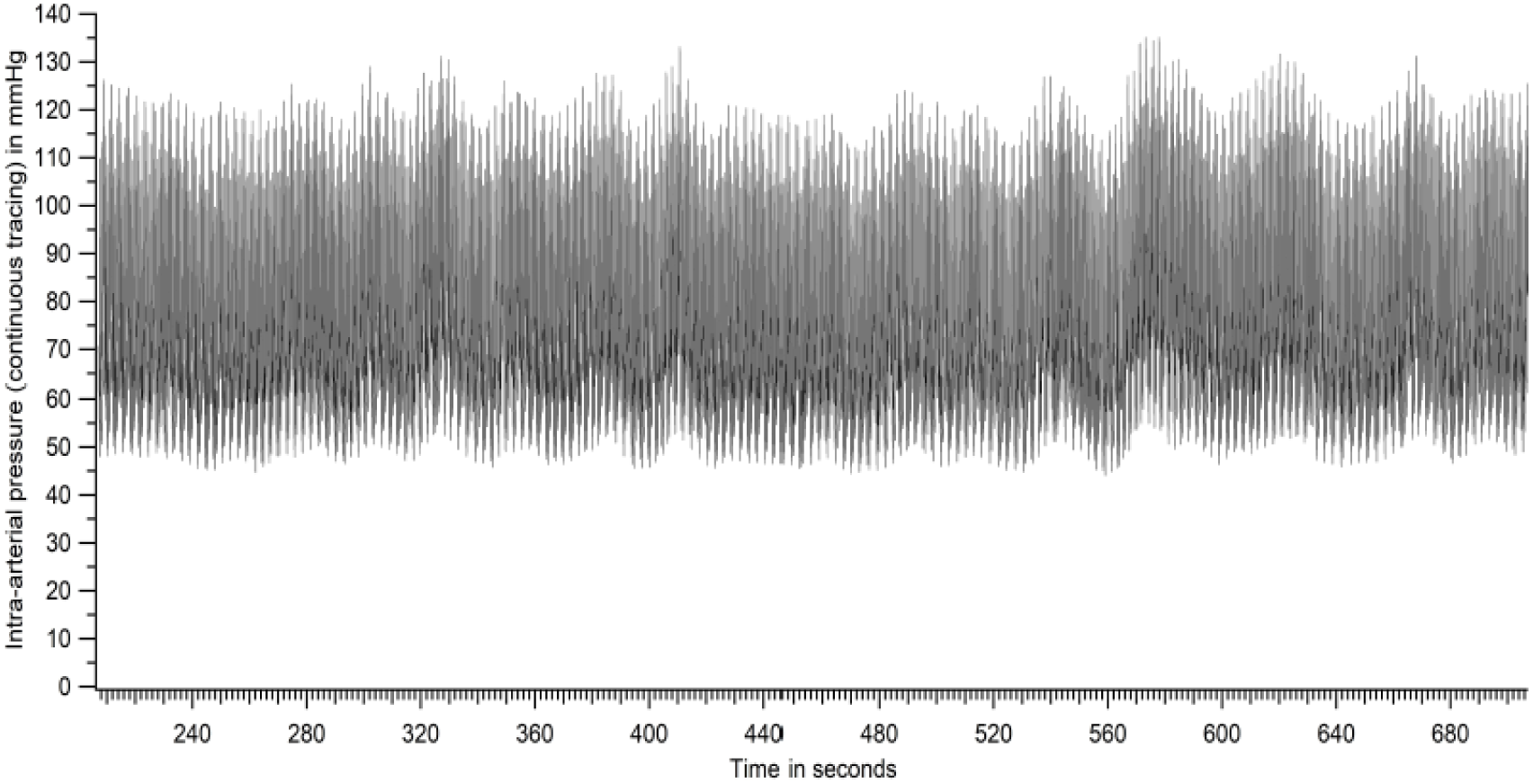
Raw tracing of intra-arterial pressure - entire recording period (patient number 46 based on Figure 17)

**Figure-5.**
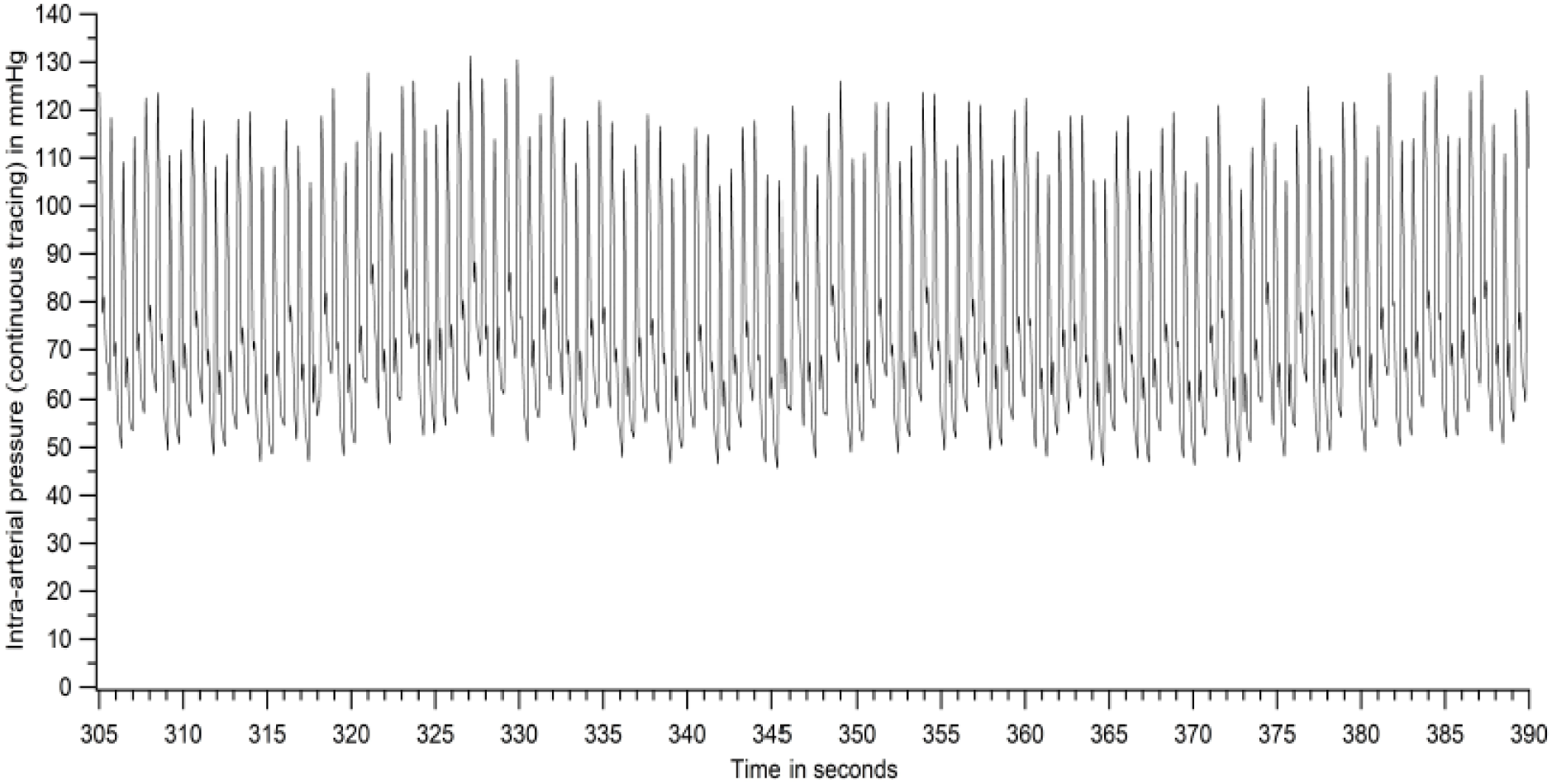
Raw tracing of intra-arterial pressure - a section of figure 4 expanded (patient no 46)

Peak (SBP) and trough (DBP) of each pressure cycle was detected using a custom-written program. Figure 6 represents the plot of systolic and diastolic pressures of each pressure cycle. Figure 6 is derived from the raw data of Figure 4.

**Figure-6.**
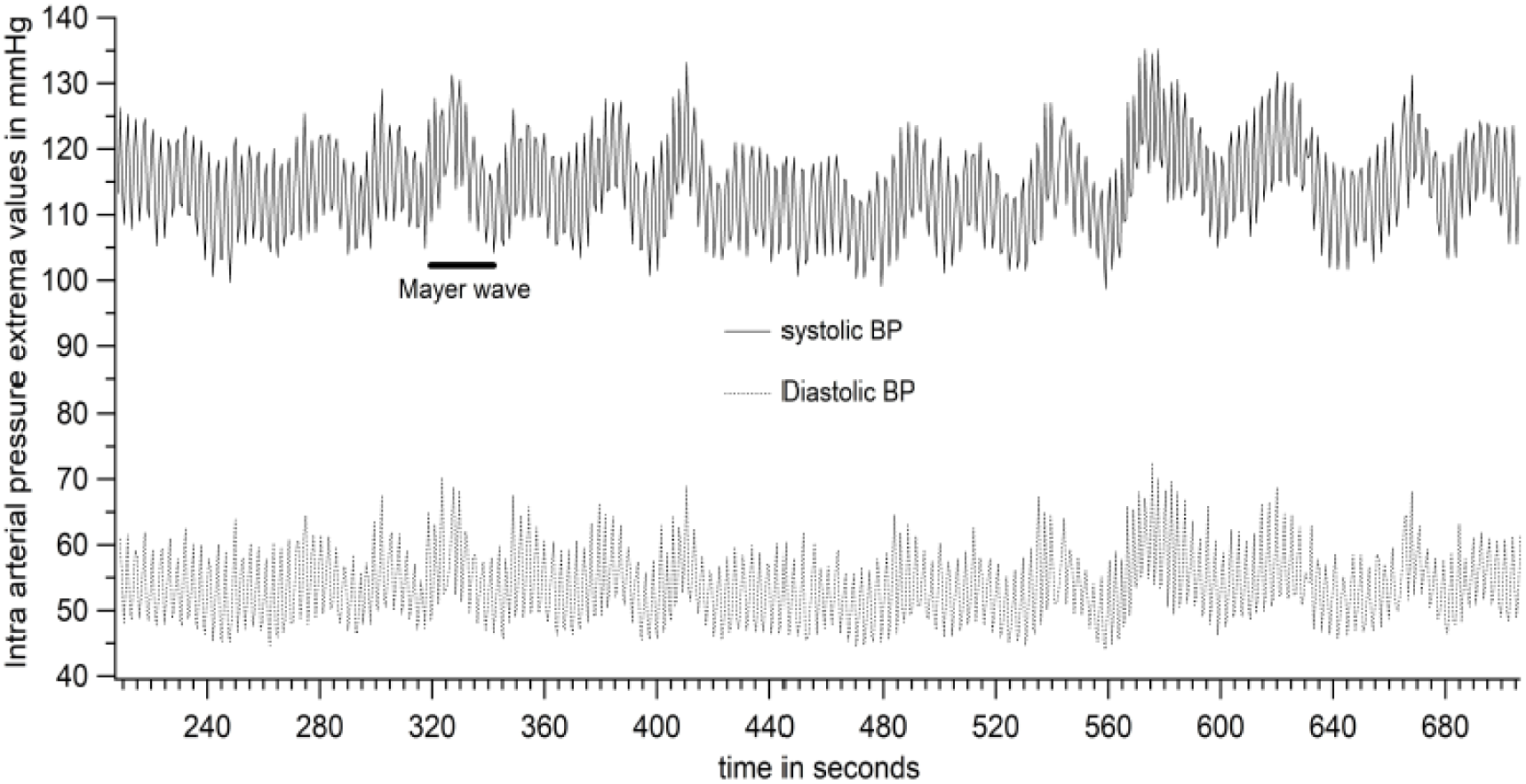
Plot of systolic and diastolic pressures of each pressure cycle of the raw tracing in Figure 4 (patient no 46)

The first order variation in arterial pressure occurs at the heart rate. Two other lower frequencies of variations are known to occur in arterial pressure. One at the respiratory frequency is referred to as Traube waves (second order variation), and another at a frequency lower than respiratory frequency (third order variation) is referred to as Mayer waves.

Figure 7 is an expansion of a section of Figure 6 and the data points for every pressure cycle are also shown. The Traube and Mayer waves are clearly seen in figures 6 and 7. Peaks and troughs of the systolic pressure variation in figure 6 (maximum and minimum systolic pressures of a Traube wave respectively) as well as the diastolic pressure variation (maximum and minimum diastolic pressures of a Traube wave in diastolic pressures) were plotted to demonstrate the Mayer waves alone (Figure 8). As seen in figure 8, there are 4 parameters of Mayer waves, two of them representing variations in maximum and minimum systolic pressures of the systolic Traube wave and 2 others representing variations in maximum and minimum pressures in the diastolic Traube wave.

**Figure-7.**
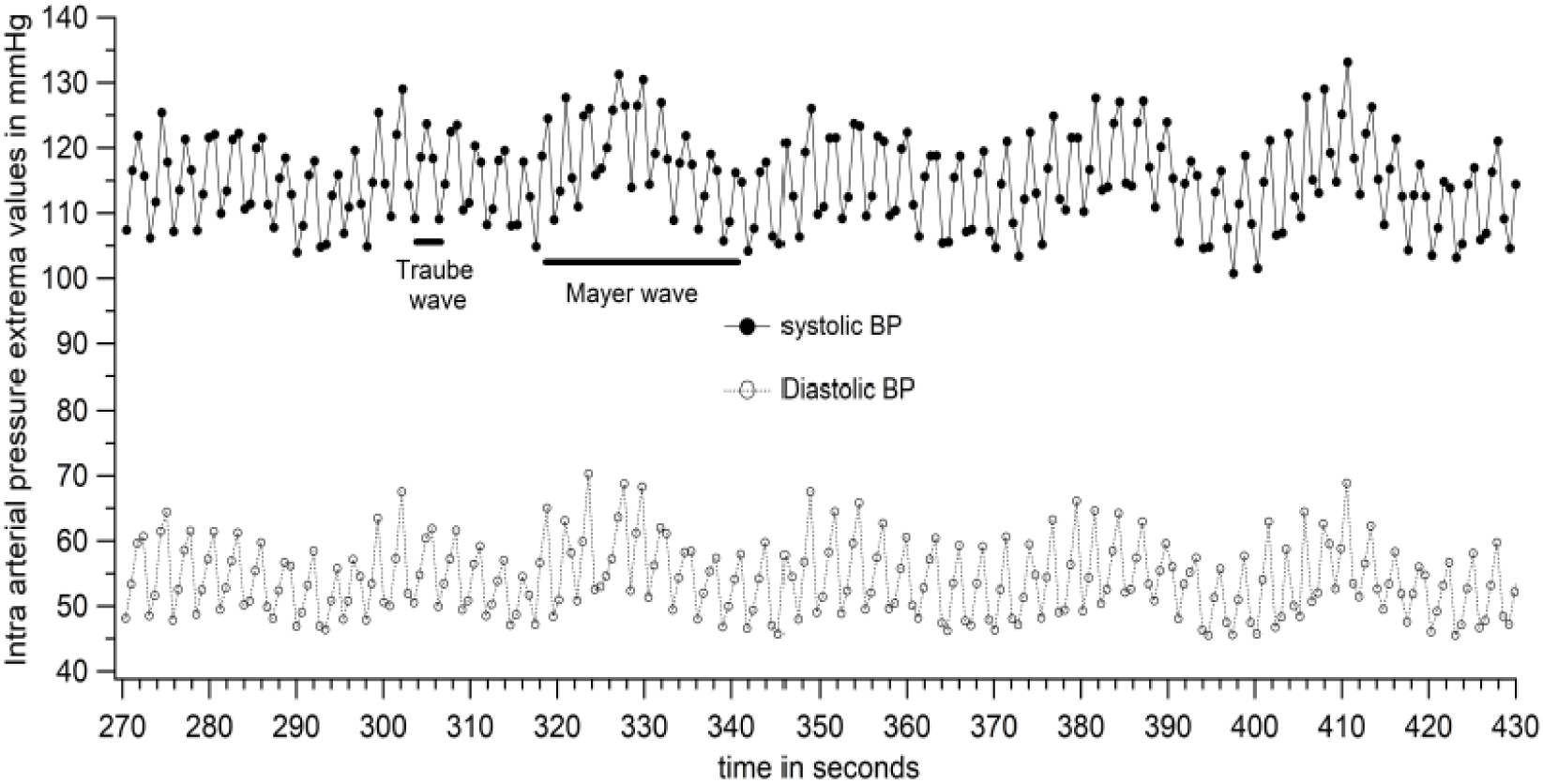
Systolic and diastolic pressure variations (expanded section from figure 6) with data points for each pressure cycle shown (patient no 46)

**Figure-8.**
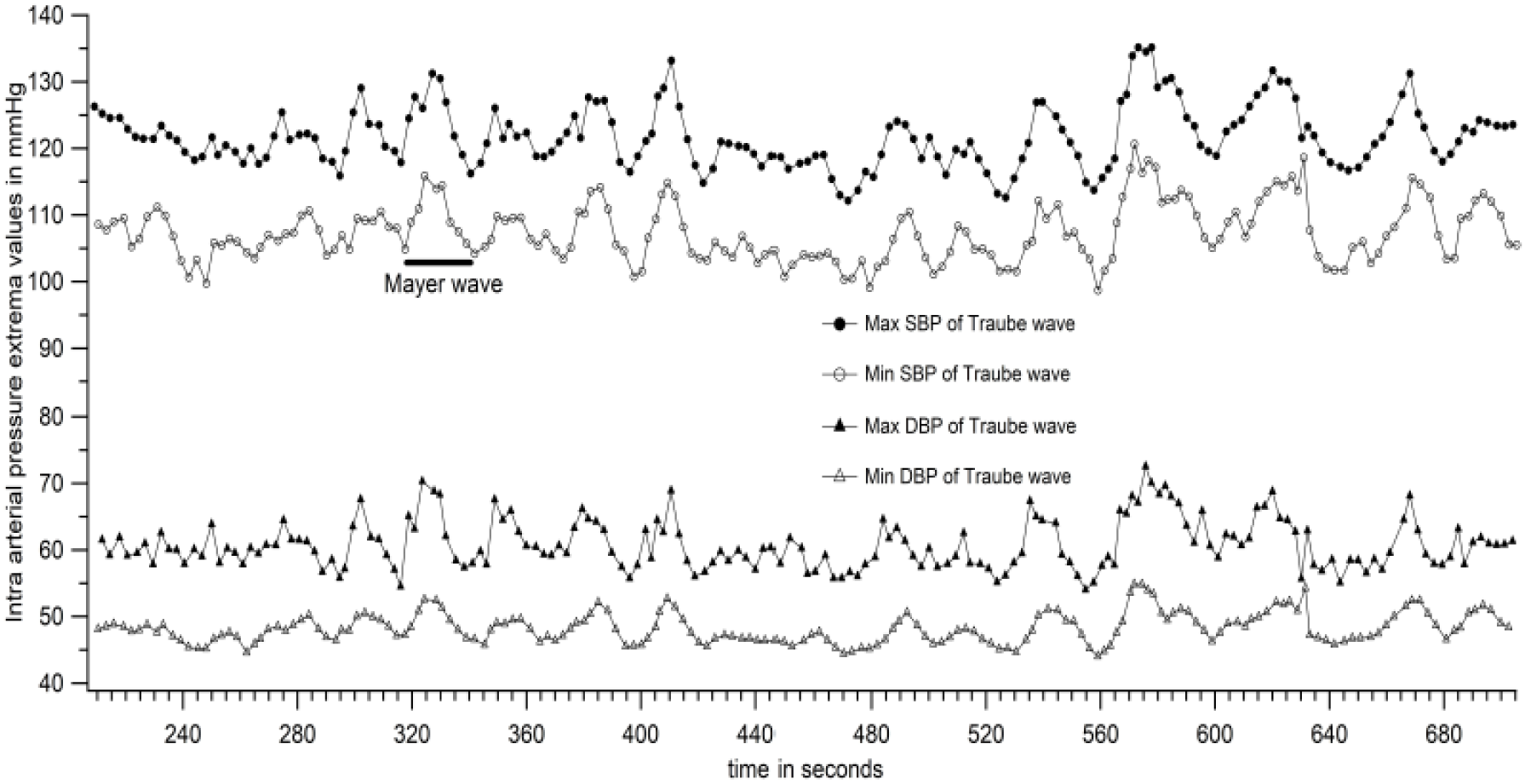
Peaks and troughs of systolic and diastolic Traube waves (seen in figures 6 and 7) plotted to get 4 Mayer wave components (patient no 46)

The Power Spectrum of data in figure 4, was calculated with the Discrete Fourier Transform (using an FFT or Fast Fourier Transform algorithm) with frequency resolution of 0.0076 Hz (about 700 seconds of data with sampling rate of 1000 Hz, average of 100 overlapping FFT periodograms, each data block zero padded to 131072 points, Hamming windowed) and is shown in figure 9. The most prominent peaks observed were the one at the heart rate, and higher harmonics of that frequency. Two well-separated peaks below the heart rate were observed in most patients. These are the frequencies corresponding to the Traube and Mayer waves.

**Figure-9.**
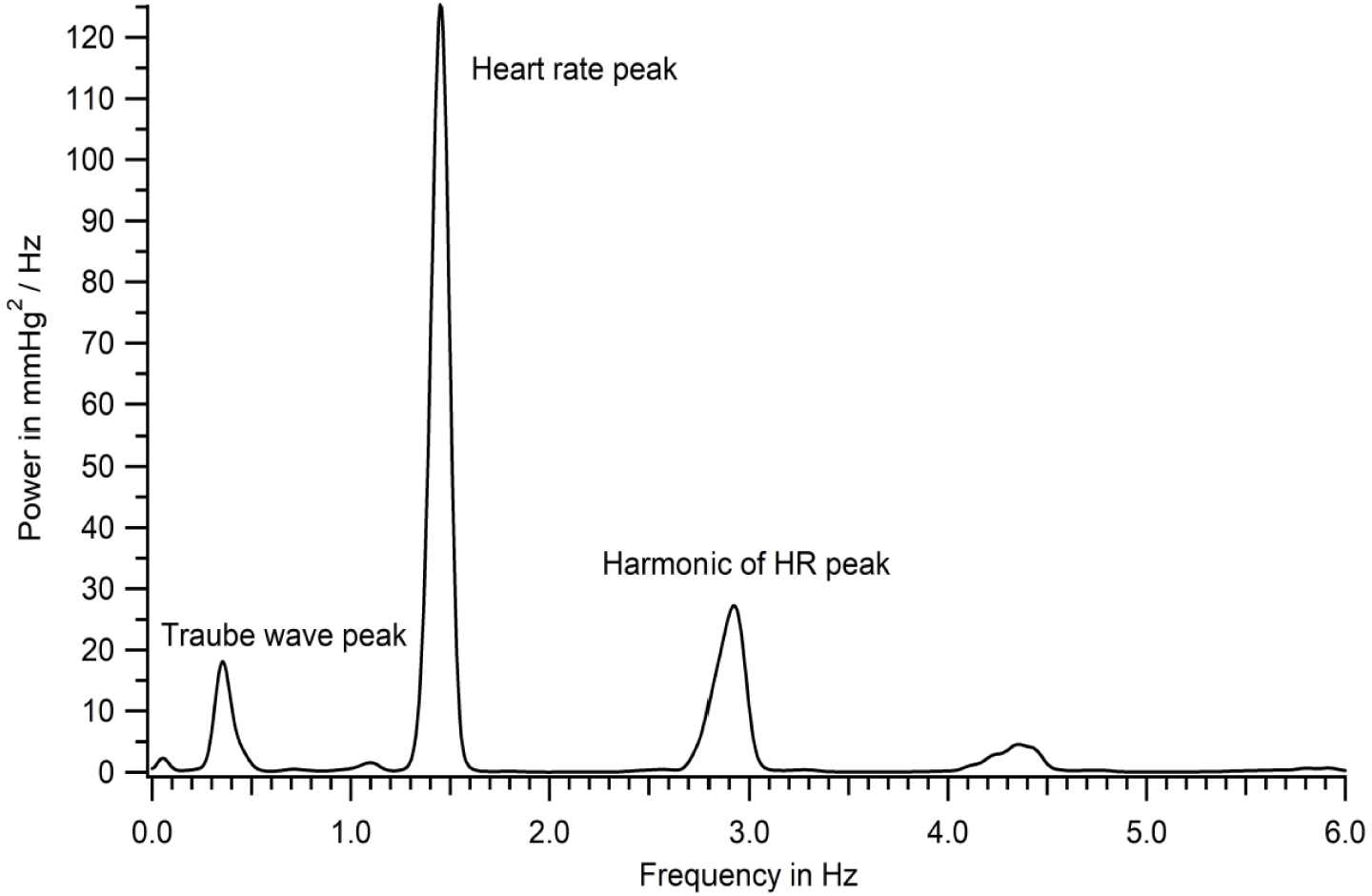
Power spectrum of the intra-arterial recording shown in figure 4 (patient no.46)

### Criteria for identifying Mayer and Traube waves from DFT spectrum

The lowest frequency peak in the DFT spectrum, which was also less than 0.12 Hz is identified as Mayer wave peak. When there were only two peaks to the left of the heart rate peak in the DFT, the higher one was taken as the Traube wave peak; when there were multiple peaks to the left of the heart rate peak, the one that was closest to the Traube wave frequency calculated by a second method (as inverse of wave period (see row 6 of Table 1)) was taken as the Traube wave peak.

**Table 1:** Mean pressure, variations in pressure and frequency of Traube waves for the recording shown in figures 4 and 6 (patient number 46) for the entire duration of recording. All values given as mean ± SD.

|  | Systolic | Diastolic |
| --- | --- | --- |
| Mean Pressure (mm Hg) | 114 ± 7 | 54 ± 6 |
| Beat to beat variation (mm Hg) | 7 ± 4 | 6 ± 4 |
| Traube wave amplitude (mm Hg) | 14.5 ± 2.5 | 13 ± 3 |
| Range of pressure (mm Hg) (minimum to maximum) | 100 – 133 | 45 – 69 |
| Total variation (mm Hg) (Difference between maximum and minimum pressures) | 33 | 24 |
| Traube wave frequency (Hz) computed as inverse of wave period | 0.37 ± 0.07 | 0.37 ± 0.07 |

Figure 10 is an expansion of Figure 9 to show clearly the dominant frequency of Traube (0.354 Hz) and Mayer waves (0.054 Hz) for the recording shown in figures 4-8. The height of Mayer waves in the spectrum is much smaller than that of Traube waves in Figures 9 and 10. However, this was not consistent. In some patients, the Mayer wave amplitude was larger. One such power spectrum is presented in Figure 11.

**Figure-10.**
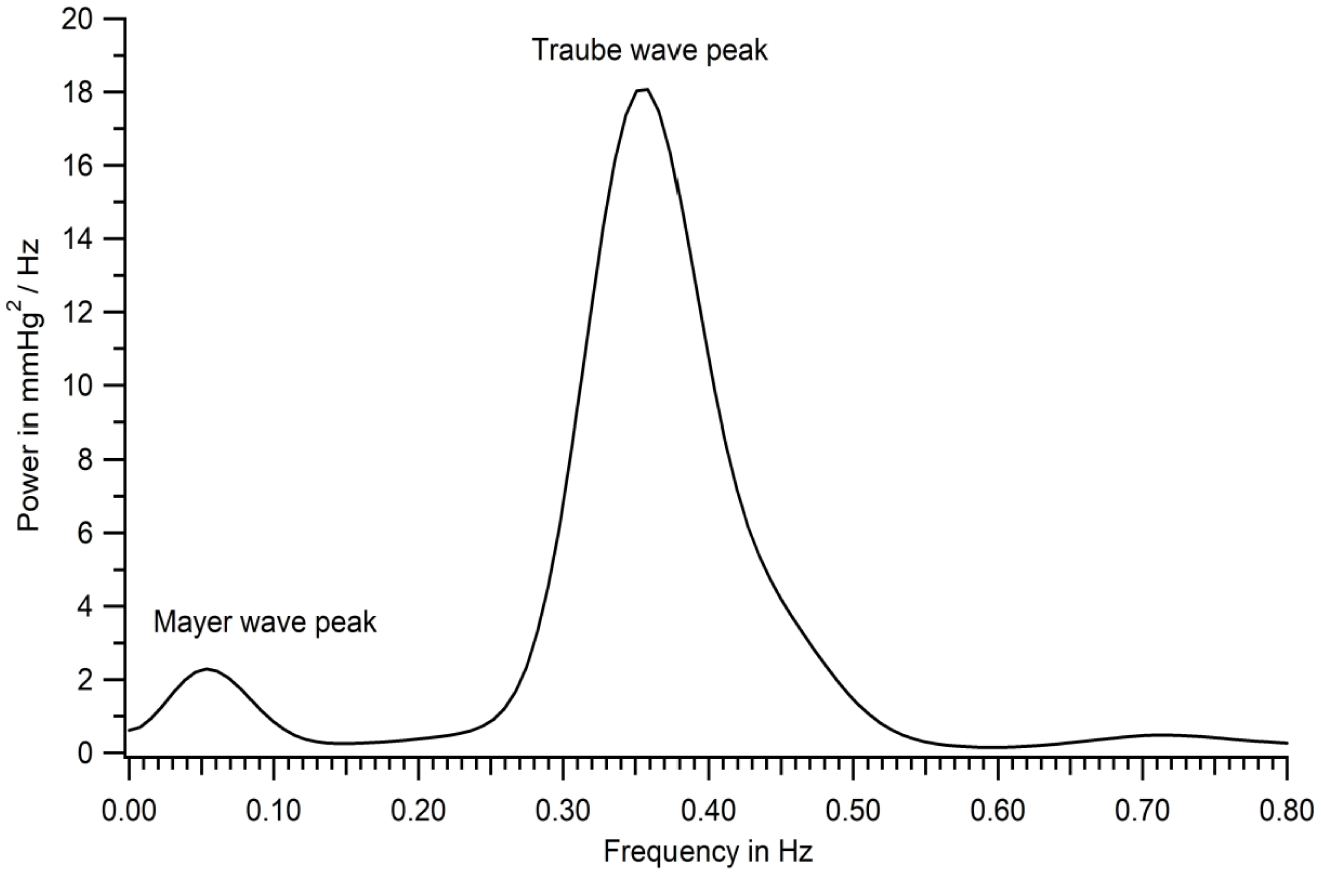
Power spectrum shown in figure 9 expanded to demonstrate Mayer wave peak (patient no 46)

**Figure-11.**
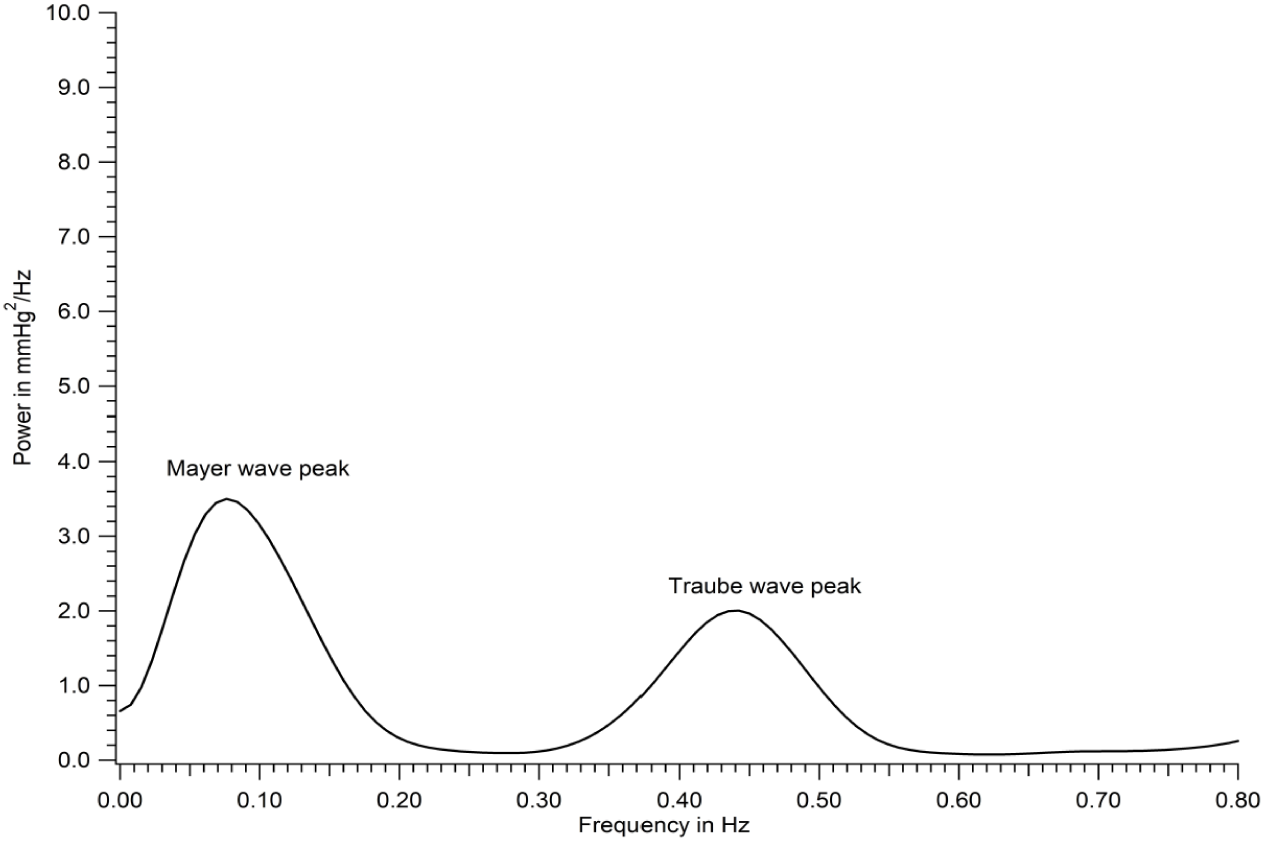
Power spectrum where Mayer peak was higher (patient number 41 of figure 17)

### Variations in pressure

Systolic pressure from every pressure cycle during the recording period (in figure 4) were averaged to get the **mean systolic pressure** in the individual during that period. Standard deviation of systolic pressure from the mean was also determined. These data are shown in row 1 of table 1.

**Beat to beat variation** in systolic pressure was calculated as the difference between two consecutive systolic pressure values. The values for the whole recording were averaged and mean ± SD of beat to beat variation is shown in row 2 of table 1.

**Traube wave amplitudes** were computed for every wave seen in figure 6 as the difference between peak and trough of the Traube waves. Amplitude of Traube waves is reported as mean ± SD for the duration of the recording in row 3 of table 1.

**Maximum systolic pressure** during the recording was determined by averaging 10 highest values of maxima of the Traube waves seen in figure 6. **Minimum systolic pressure** was similarly determined by averaging 10 lowest values of minima of systolic Traube waves. This was done to avoid overestimations of variation due to a momentary large deflection of pressures.

Data for maximum and minimum systolic pressures in the individual is shown in row 4 of table 1. **Total magnitude of systolic pressure variation** in the individual is calculated as the difference between maximum and minimum systolic pressures obtained as just stated. The total variation in systolic pressure during the recording period is given in row 5 of table 1.

**Traube wave frequency** was computed as inverse of wave period for every Traube wave cycle. Mean and SD of the frequency was calculated for the entire duration of the recording and the data is shown in row 6 of table 1.

Similar calculations were done for diastolic pressures and the data is found in table 1.

Data in rows 1 and 4 of Table 1 is presented in figure 12. Data for systolic pressure variability in rows 2, 3 and 5 are presented in figure 13 and that for diastolic pressure variability in figure 14.

**Figure-12:**
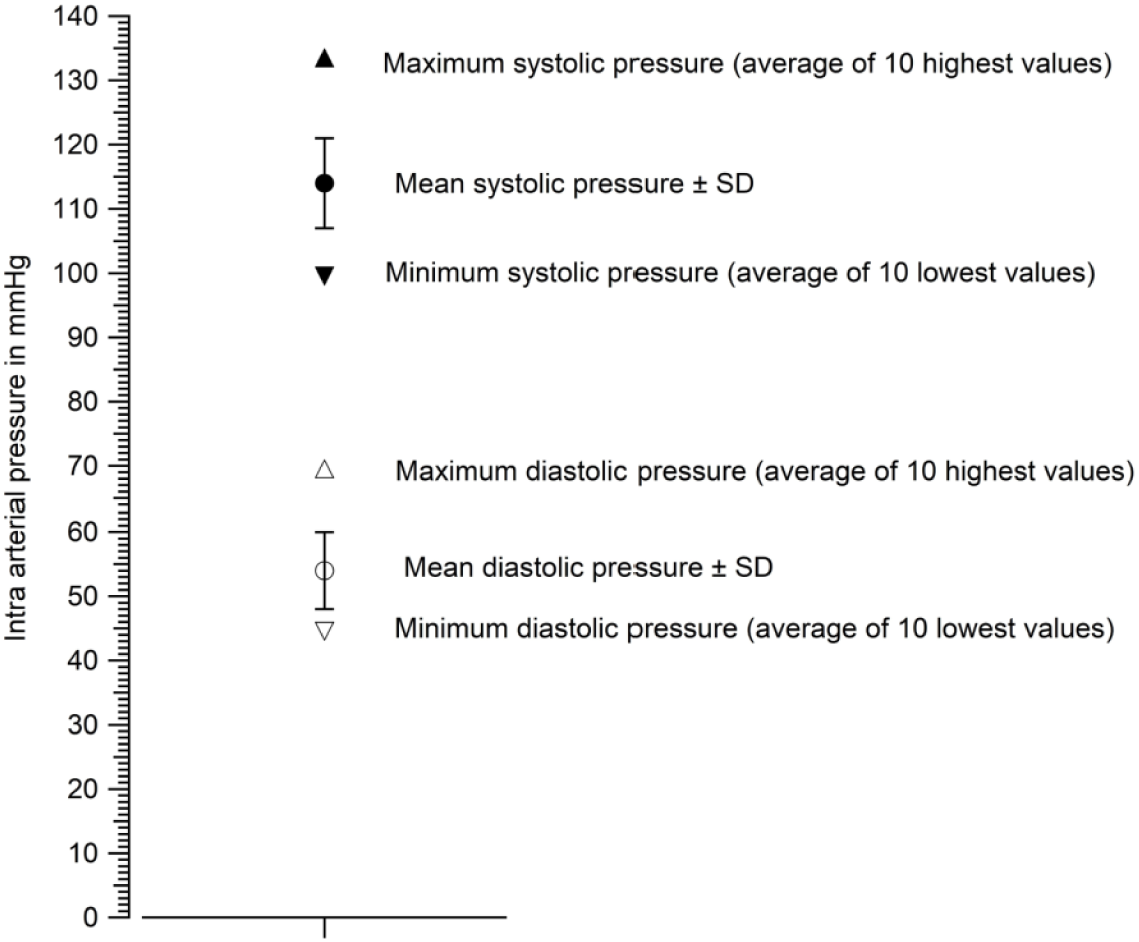
Mean (± SD), maximum and minimum systolic and diastolic pressures for the tracing shown in figure 4 (from table 1, (patient no 46))

**Figure-13.**
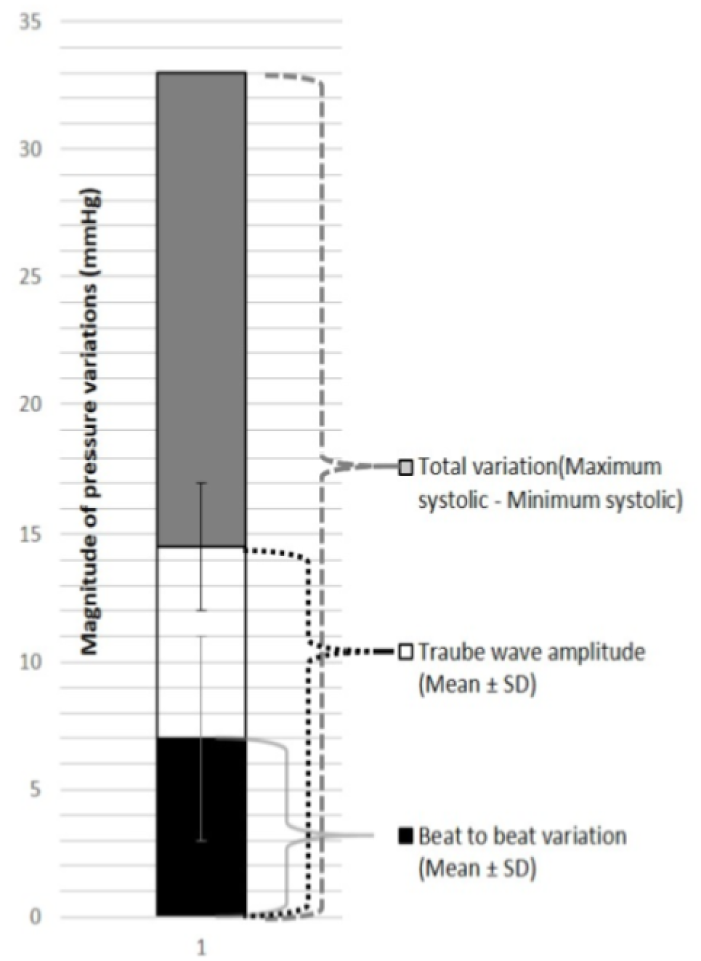
Systolic pressure variability for the recording in Figure 4 (patient no 46)

**Figure-14.**
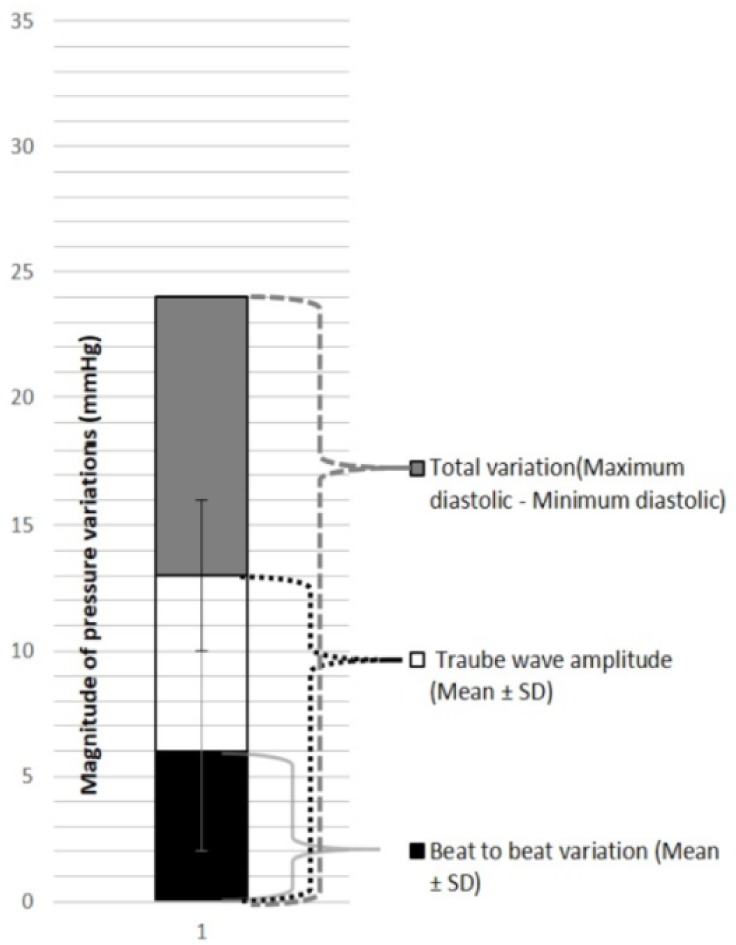
Diastolic pressure variability for the recording in Figure 4 (patient no 46)

Data from all recordings were subjected to analyses as above and the data for all 51 patients is presented in the following sections.

### Data for the whole sample

#### Frequency of BPV in the study sample

Discrete Fourier transform (DFT) spectrum calculation were performed on each intra-arterial pressure recording (see figure 15). Peak frequencies of Traube and Mayer waves were obtained from DFT power spectra. Frequency of Traube waves was also calculated as the inverse of duration of each Traube wave and averaged. A plot of the frequencies obtained for all 51 patients is shown in figure 15. In 49 out of 51 patients there were clear peaks on the DFT spectrum that could be identified as Mayer wave peaks as per our criteria. Figure 16 is a histogram of Mayer wave frequencies in the study sample. From figure 16 and 17, it can be stated that the major Mayer wave frequency band in the study sample was 0.045 to 0.065 Hz.

**Figure-15.**
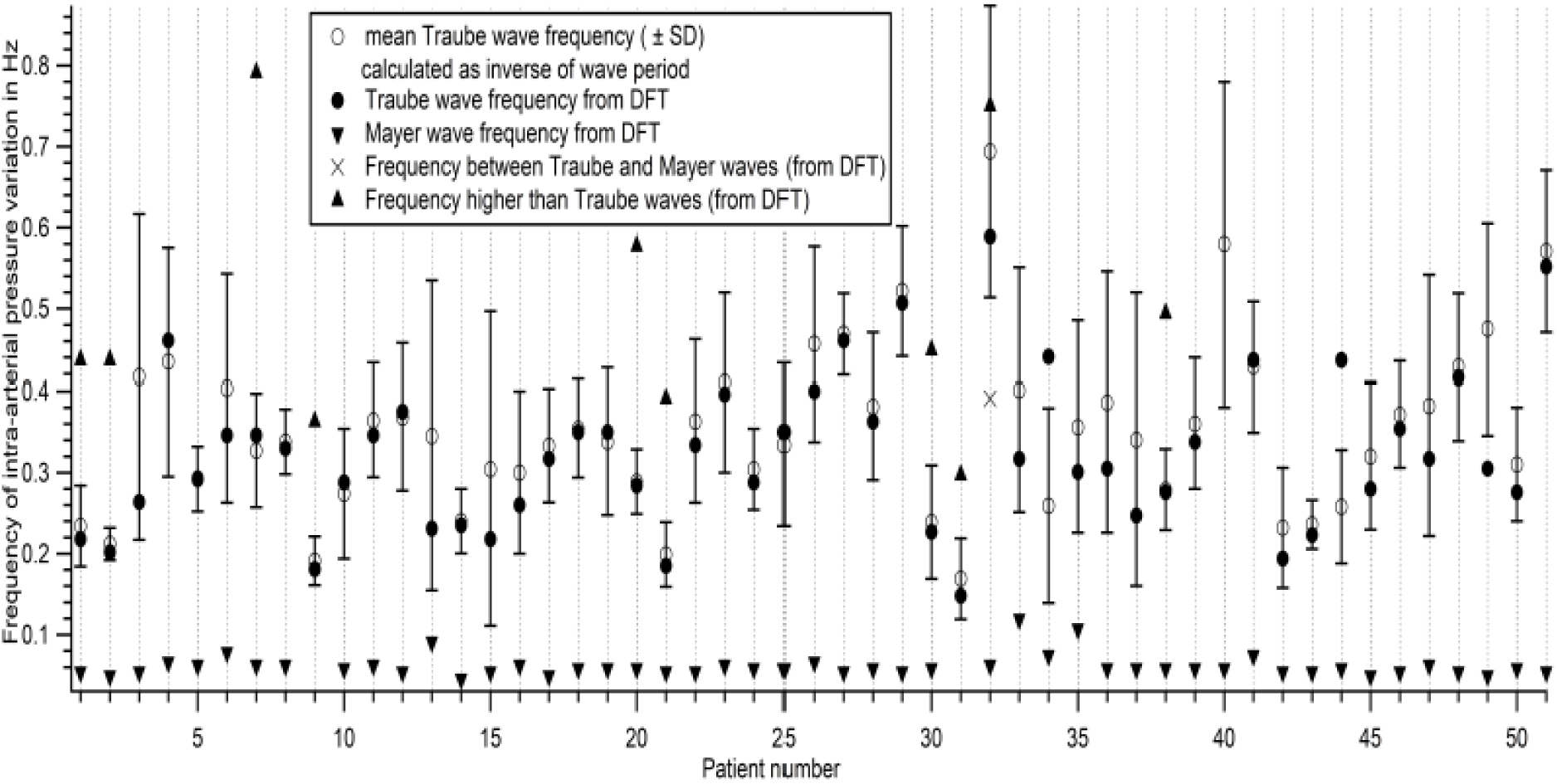
Frequencies of pressure variations in 51 patients

**Figure-16.**
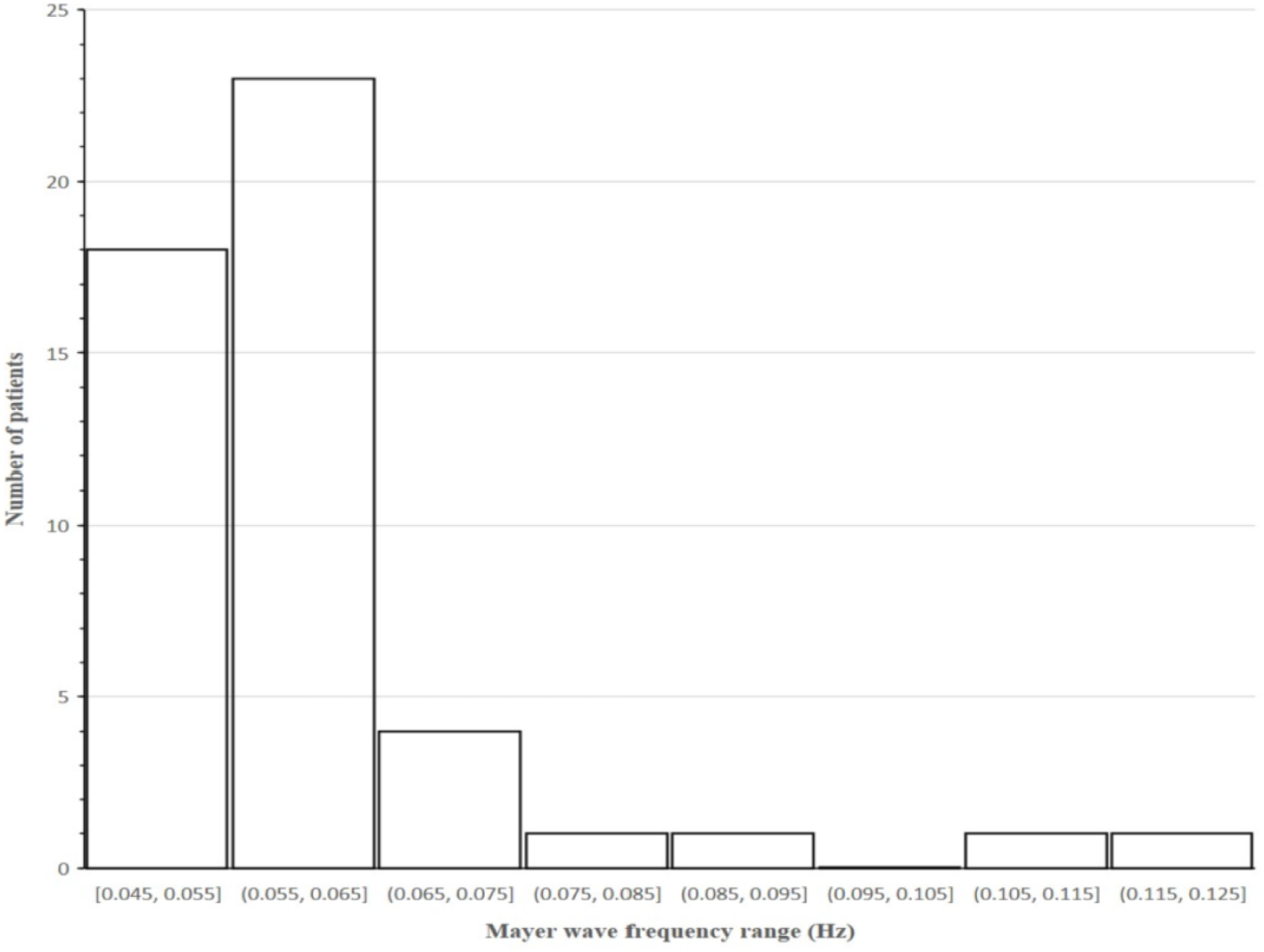
Histogram of Mayer wave frequency in the sample (49/51 patients)

**Figure-17.**
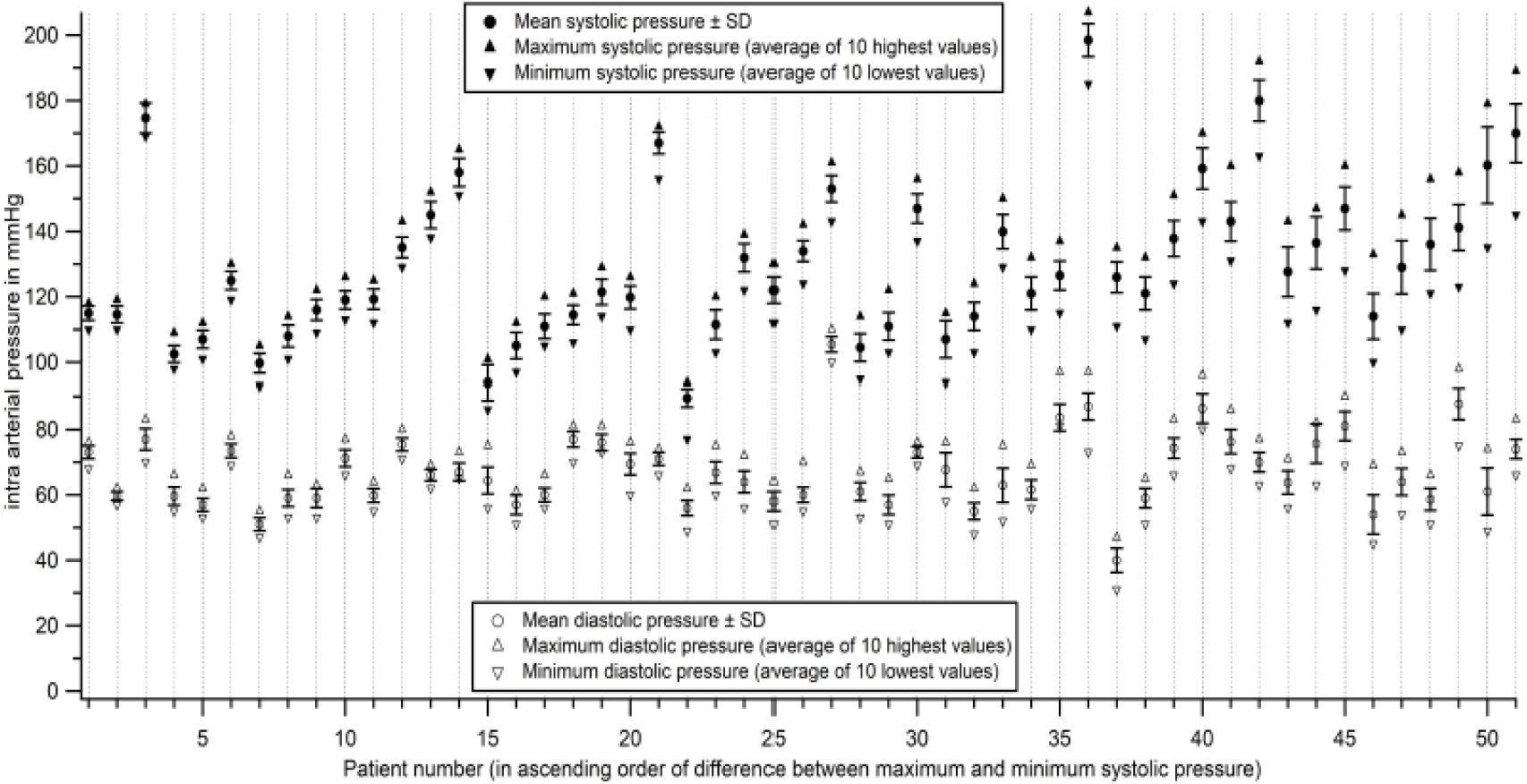
Mean (± SD), minimum and maximum Systolic and diastolic pressures of the 51 patients

The parameters found in Table 1 (for a single patient) were calculated for all 51 patients. Magnitude of variations in systolic and diastolic pressures (as represented for one patient in figure 12-14) is shown in figures 17 and 18 for all 51 patients.

**Figure-18.**
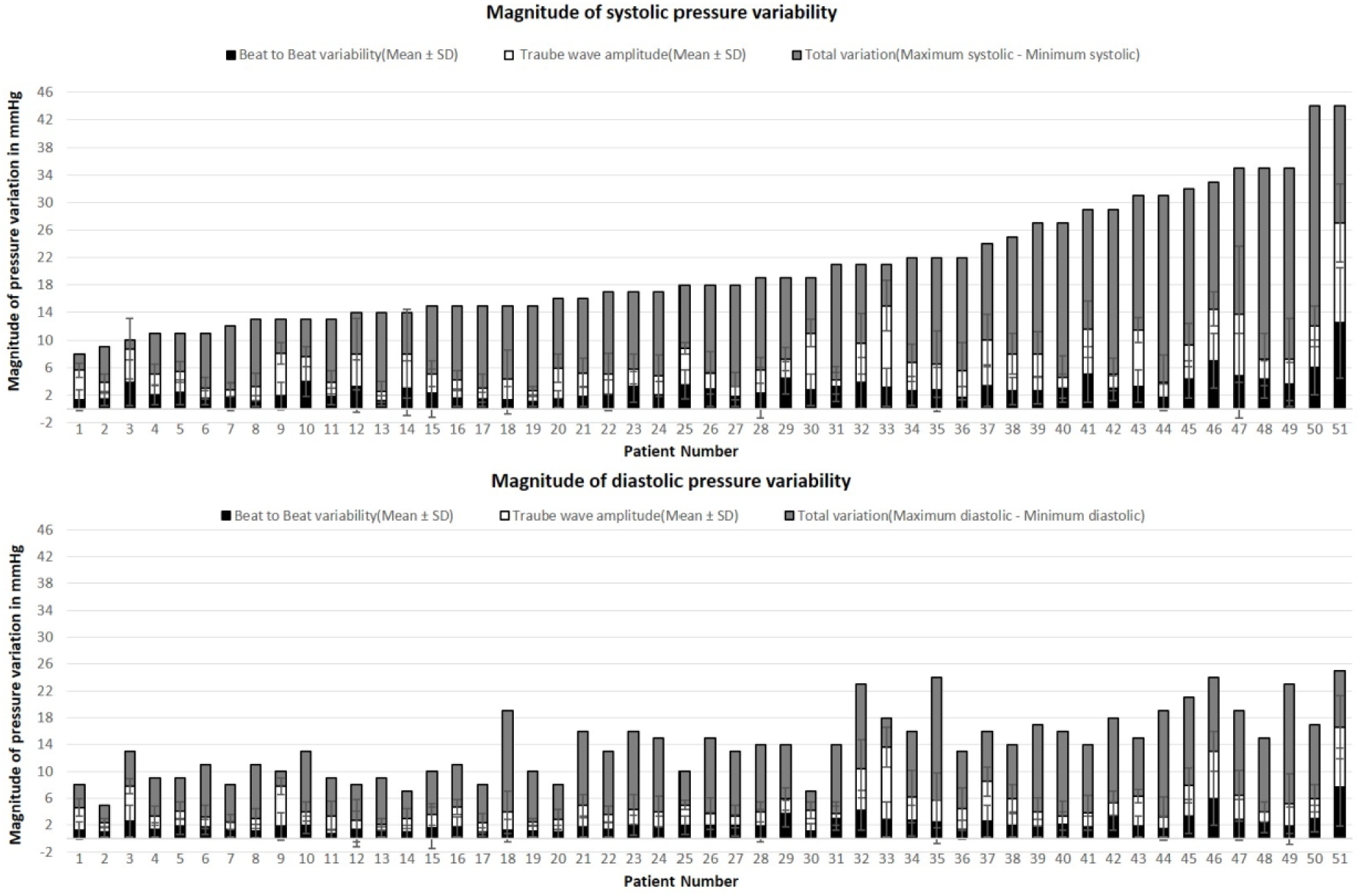
Beat to beat variability, Traube wave amplitude and total magnitude of variation in systolic and diastolic pressures in 51 patients (data arranged in ascending order of total systolic variation and hence the patient numbers)

Sample means were calculated for all parameters in Table 1 and the values are given in Table 2. The Mean value (± SD) for the magnitude of systolic pressure variation in the sample was determined to be 21 ± 9 mmHg and that of diastolic pressure variation as 14 ± 5 mmHg.

**Table 2:** Sample mean (SD) of parameters for every patient as found in Table 1 (Values for pressures rounded to the nearest whole number).

|  | Systolic | Diastolic |
| --- | --- | --- |
| Mean of Mean Pressure(mmHg) | 130 (23) | 67 (11) |
| Mean Beat to beat variation(mmHg) | 3 (2) | 2 (1) |
| Mean Traube wave amplitude (mmHg) | 7 (4) | 5 (3) |
| Mean range of pressure (mmHg) i.e., mean minimum pressure to mean maximum pressure | 119 (22) - 130 (25) | 60 (11) – 74 (12) |
| Mean Total variation (mm Hg) (mean difference between maximum and minimum pressures) | 21 (9) | 14 (5) |
| Mean Traube wave frequency (Hz) (computed as inverse of wave period) | 0.35 (0.1) | 0.35 (0.1) |

## Discussion

### Dynamic characterization of the catheter-transducer system

The dynamic response of the fluid-filled catheter-transducer system must be sufficiently good to reproduce the pressure waveform with high fidelity and minimal distortion. If the natural frequency of the catheter system overlaps the frequency band of the signal, there will be resonance and undue amplification of the signal at the resonant frequency. Therefore, the catheter system must have a natural frequency much higher than the highest frequency component of the recorded signal. In the case of arterial pressure, the highest frequency component of the pressure pulse would be the fast upstroke during systole.

In addition to the natural frequency of the catheter system, the other parameter that determines the fidelity of the recorded signal is the damping coefficient. Under-damping can lead to falsely high systolic pressures and resonant ringing of the pressure pulse [2], while over-damping will conversely lead to diminution of the amplitude, loss of details like the dicrotic notch in addition to falsely low systolic pressures.

Natural frequency and damping co-efficient are not quantified in current clinical practice. The assessment of dynamic characteristics is qualitative based on display of the fast-flush test on the monitor. Small changes in natural frequency and damping co-efficient can significantly alter wave form, resulting not only in errors in systolic and diastolic pressures but also in timing of systolic peak and dicrotic notch, and these errors will propagate in calculation of other parameters, for e.g., pulse transit time (PTT). For accuracy of pressure measurement, quantitation of the natural frequency and damping co-efficient must become part of routine clinical practice.

#### Frequency of Blood pressure variations

An important observation in this study is the frequency band of the Mayer waves or the third order variations in blood pressure. Traube was the first to study variations in blood pressure and observed the respiratory (second order) variations and third order variations occurring at a frequency lower than respiratory variations. Mayer is reported to have studied the slower third order variations further [11]. The third order variations are commonly referred to as Mayer waves by many research groups [12-15] though there are references to these as Traube-Hering-Mayer waves and even arguments that they must be referred to as Traube waves only after the original discoverer [16]. There are others who have referred to the respiratory variations (second order) as Traube waves and the low frequency variation (about 0.1 Hz) as Mayer waves [17]. In keeping with current trends, we have referred to the low frequency variation (third order) as Mayer waves and to give credit to the pioneer in the field, while avoiding confusion, we have referred to the respiratory variation in arterial pressure as Traube waves, along the lines of Karemaker (1999) [17].

The Mayer wave frequency has been reported to be about 0.1 Hz in several articles [15,18] while others have given a range of 0.05 to 0.15 Hz [14,19]. In this context, we draw attention to the fact that in 49 out of 51 patients who had a Mayer wave peak in the DFT, the frequency was less than 0.12 Hz. In 41 out of 51 patients, the Mayer wave frequency range was 0.045 Hz to 0.065 Hz. It may therefore be stated that the major Mayer wave frequency band in humans may be 0.045 to 0.065 Hz, keeping in mind that the study sample is a patient population, however, haemodynamically stable. The other possibility is that Mayer wave frequency shifts to a lower range in acute illness.

The Mean value (± SD) for the magnitude of systolic pressure variation in the sample was determined to be 21 ± 9 mm Hg and that of diastolic pressure variation as 14 ± 5 mm Hg. These values are not from a normal population and may not represent normal variability. What we intend to convey is that while the systolic pressure in an individual can vary by as much as 21 mm Hg or more during a few minutes of recording, getting a point value for systolic pressure as provided by either the manual Korotkoff method or oscillometric method can lead to misinterpretations about the cardiovascular status of the individual. For instance, in patient number 51 of figures 17 and 18, systolic pressure varies between 145 and 189 mm Hg. The current non-invasive methods, if otherwise accurate, will report a single point value anywhere along this range of 145 mm Hg to 189 mm Hg for this patient.

The latest American and European guidelines for diagnosis and treatment of hypertension depend on single point cut-offs for systolic as well as diastolic pressures [20]. However as documented by others and as shown here, systolic and diastolic pressures vary considerably even over a few seconds and a blood pressure report should ideally contain the range of systolic and diastolic pressures in the individual during the recording period. In the study sample, there are 14 patients whose systolic pressure range includes 140 mm Hg which is the cut=off for diagnosing hypertension. Single point estimates of systolic pressure in these individuals will yield values anywhere in the range, i.e., higher than or lower than 140 mm Hg (if there are no measurement errors) and probabilistically, the values will differ with every measurement making it difficult to decide whether to treat or not.

In figure 17, it is also noted that the Traube wave amplitude in most patients is less than half of the total variability. This is true even in patients in whom the power of the Mayer wave in a DFT spectrum is much less than that of the Traube wave, where the difference between the total variation (including the Traube wave amplitude) and the Traube wave amplitude is higher than the Traube wave amplitude *per se* (see figures 10 and 13- they are from patient number 46). We may infer from this that the magnitude of the overall variation even over a period of 10 minutes is not well-captured by estimation of the Traube wave amplitude – there are other variations that may be irregular and consequently of lower frequencies.

## Conclusion

The magnitude of systolic and diastolic pressure variations in a period as short as 10 minutes is sizeable and therefore it is imperative that methods of assessing the entire range of pressures in an individual are developed for meaningful assessment of cardiovascular status.

## Acknowledgements

The Authors are grateful to the medical and technical personnel at the Surgical Intensive Care Unit of Christian Medical College, Vellore for enabling the blood pressure recordings. Mrs. Surekha (Junior Research Fellow, DBT) is acknowledged for technical help in analysis of data and manuscript preparation.

## References

1. Parati G, Stergiou GS, Dolan E, Bilo G. Blood pressure variability: clinical relevance and application. The Journal of Clinical Hypertension. 2018; 20(7) : 1133–1137.

2. Romagnoli S, Ricci Z, Quattrone D, Tofani L, Tujjar O, Villa G, Romano SM, Gaudio AR. Accuracy of invasive arterial pressure monitoring in cardiovascular patients: an observational study. Critical Care. 2014; 18(6):644.

3. Gibbs NM, Larach DR, Derr JA. The Accuracy of Finapres™ Non-invasive Mean Arterial Pressure Measurements in Anesthetized Patients. Anesthesiology. 1991;74(4):647–652.

4. Silke B, Spiers JP, Boyd S, Graham E, McParland G, Scott ME. Evaluation of non-invasive blood pressure measurement by the Finapres method at rest and during dynamic exercise in subjects with cardiovascular insufficiency. Clinical Autonomic Research. 1994; 4(1-2):49–56.

5. Agnes S. Meidert, Saugel B. Techniques for Non-Invasive Monitoring of Arterial Blood Pressure. Frontiers in Medicine. 2018; 4:231.

6. Parati G. Blood pressure variability, target organ damage and antihypertensive treatment. Journal of Hypertension. 2003; 21(10) :1827–1830.

7. Tatasciore A, Renda G, Zimarino M, Soccio M, Bilo G, Parati G, Schillaci G, De Caterina R. Awake systolic blood pressure variability correlates with target-organ damage in hypertensive. Hypertension. 2007; 50(2):325–332.

8. Leoncini G, Viazzi F, Storace G, Deferrari G, Pontremoli R. Blood pressure variability and multiple organ damage in primary hypertension. Journal of Human Hypertension. 2013; 27:663–670.

9. Rothwell PM. Does Blood Pressure Variability Modulate Cardiovascular Risk? Current Hypertension Reports. 2011;13(3):177–186.

10. Gardner RM. Direct blood pressure measurement--dynamic response requirements. Anaesthesiology. 1981;54:227–236.

11. Wood JR, HC. The origin of the “Traube” waves. American Journal of Physiology-Legacy Content. 1899;2(4):352–354.

12. Killip T. Oscillation of blood flow and vascular resistance during Mayer waves. Circulation research. 1962; 9: 987–993.

13. Elghozi JL, Laude D, Girard A. Effects of respiration on blood pressure and heart rate variability in humans. Clin.exp.pharmacol.physiol. 1991;18(11):735–742.

14. Takalo R, Korhonen I, Majahalme S, Tuomisto M, Turjanmaa V. Circadian profile of lowfrequency oscillations in blood pressure and heart rate in hypertension. American Journal of Hypertension. 1999; 12 (9):874–881.

15. Julien C. The enigma of Mayer waves: Facts and models. Cardiovascular Research. 2006; 70:12–21.

16. Halliburton WD. Traube waves and Mayer waves. Quarterly Journal of Experimental Physiology. 1919; 12(3):227–229.

17. Karemaker JM. Autonomic integration: the physiological basis of cardiovascular variability. The Journal of Physiology. 1999; 517 (Pt 2) 316.

18. Hamner JW, Morin RJ, Rudolph JL, Taylor JA. Inconsistent link between low-frequency oscillations: R-R interval responses to augmented Mayer waves. Journal of Applied Physiology. 2001; 90(4):1559–1564.

19. Taylor JA, Williams TD, Seals DR, Davy KP. Low-frequency arterial pressure fluctuations do not reflect sympathetic outflow: gender and age differences. Am J Physiol. 1998;274:H1194–1201.

20. Harrap et al. New Blood Pressure Guidelines Pose Difficult Choices for Australian Physicians. Circulation Research. 2019;124:975–977.

